# AI-powered integration of multi-source data for TAA discovery to accelerate ADC and TCE drug development (I): TAA Target Identification and Prioritization

**DOI:** 10.1101/2025.05.06.652559

**Authors:** Tao Xie, Chao-Hui Huang

## Abstract

The advancement of T-cell engagers (TCEs) and antibody-drug conjugates (ADCs) has been hindered by fragmented data landscapes. This paper, the first in a series, introduces an AI-driven framework specifically for tumor-associated antigen (TAA) target identification and prioritization, a critical initial step in TCE and ADC development. Our framework integrates diverse datasets— including multi-omics repositories and information from scientific publications—to systematically enhance the discovery of TAAs. We have developed a graph retrieval-augmented generation (RAG)-enhanced language model that extracts insights from biological and clinical literature, while integrating curated public oncology-related omics databases such as TCGA, GTEx, single-cell atlases, and additional omics datasets. This approach prioritizes TAAs with high tumor selectivity and low on-target/off-tumor risk. By unifying diverse knowledge sources, our method provides a scalable, efficient, and data-agnostic strategy to address attrition challenges in both ADC and TCE drug development pipelines, focusing initially on TAA target identification and prioritization to transform the landscape of cancer therapeutics.

## Introduction

Cancer remains a significant global health challenge, emphasizing the need for innovative therapeutic strategies. Tumor-associated antigens (TAAs) play a crucial role in cancer treatment by serving as precise targets for Antibody-drug conjugates (ADCs) and T-cell engagers (TCEs) [1]. These platforms are designed to eradicate cancer cells while minimizing harm to healthy tissues. TCEs function by redirecting T-cells to attack cancer cells, utilizing the body’s immune system to effectively eliminate malignant cells while ADCs leverage the specificity of antibodies to deliver potent cytotoxic drugs directly to cancer cells, ensuring targeted treatment.

Despite advancements in genomics and bioinformatics that have deepened our understanding of tumor biology, converting this knowledge into practical therapeutic TAA targets remains challenging. The success of TCEs and ADCs heavily depends on rigorous target selection. Identifying and prioritizing suitable TAAs is vital for enhancing therapeutic precision and reducing off-target effects. Without effective target triage, the ability of TCEs and ADCs to deliver focused and effective cancer treatment is significantly undermined [1].

The integration of machine learning with biological expertise, particularly through large language models (LLMs), holds promise for revolutionizing the analysis of complex biological data. In the realm of oncology, LLMs show vast potential to reshape data analysis by facilitating rapid interpretations and contributing to the development of personal treatment plans tailored to individual patient needs [2-4]. Domain-specific LLMs, aided by specific Retrieval Augmented Generation (RAG), excel in tasks defined by organizational standards. These models are tailored for real-world applications, requiring a deep understanding of context, product data, corporate policies, and industry terminology. Given the advent of LLMs promises a transformative approach, offering unprecedented opportunities in, this study demonstrates the application of a graph RAG-enhanced LLM (foundation model: ChatGPT-4) to drive the discovery of TAAs, thus enhancing target selectivity and therapeutic precision for ADC and TCE drug development.

## Materials and Methods

### Knowledge Data from PubMed

Our study builds upon existing research in graph retrieval-augmented generation (RAG) and seeks to extend its application within the domain of tumor-associated antigen (TAA) discovery. Utilizing the E-utilities suite of NCBI Web APIs, we accessed the most recent open-access full-text articles focused on TAA research in lung cancer. We crafted targeted queries using keywords such as “tumor-associated antigen” and “lung cancer” to capture a comprehensive array of relevant literature. Python-based API calls were developed to retrieve PubMed IDs (PMIDs) corresponding to these queries via the **esearch** function, interfacing with the Entrez system to locate pertinent articles. Subsequently, the **efetch** function was employed to obtain detailed metadata for each PMID, including PubMed Central IDs (PMCIDs), article titles, abstracts, and more. To maintain stability and adhere to Entrez API rate limits, batch processing with pauses was implemented. All retrieved records were stored in txt format, facilitating seamless integration with data analysis tools.

### Oncology Omics Data Process

The framework is designed to integrate diverse multi-omics data types, encompassing bulk RNA expression, single-cell omics, spatial transcriptomics, and protein expression data. This comprehensive approach significantly enhances the discovery and prioritization of tumor-associated antigens (TAAs). To illustrate the integration of omics data, we present an overview of the process for handling bulk gene expression data from the TCGA and GTEx datasets. Building on the findings of a previous study [4], we utilized bulk expression data from the UCSC RNA-seq Compendium. This resource standardizes TCGA and GTEx samples from both cancerous and normal tissues by employing advanced computational techniques to eliminate batch effects. The integration of these datasets enables a robust analysis of gene expression differences between tumor and normal tissues, thereby facilitating the identification of potential TAAs. To augment the safety assessment of TAA candidates, a safety score—developed to evaluate the expression of TAAs in normal tissues relative to tumor tissues—has been incorporated into the RAG framework. This score is derived from the integrated expression data, as outlined below:

1. Z-score Calculation: For each gene, z-scores are calculated based on its expression level across different tissue types. This involves normalizing the expression data by subtracting the mean expression level and dividing by the standard deviation, thereby highlighting deviations from average expression.
2. Assessment of Normal Tissue Expression: The z-scores provide insights into how significantly a gene’s expression deviates in normal tissues compared to tumor tissues. Genes with high z-scores (**>1**) in normal tissues indicate higher expression levels relative to the average, which could suggest potential safety concerns if those genes are targeted in cancer therapy.
3. Safety Score Integration: The final score integrates these z-scores to quantify the risk associated with targeting a particular gene as a TAA. note that normal prostate tissue was excluded from the normal tissue collection, as it is not deemed critical for this analysis.

With these steps, we have generated safety scores considering the expression levels across various normal tissues to ensure that selected TAAs have minimal expression in non-cancerous tissues, thus reducing the likelihood of off-target effects. The information can be found in the supplementary (**Supplementary Table 1**).

### Construction of Biological Knowledge GraphRAG

In this study, we chose OpenAI’s GPT-4 for its real-time responsiveness and extensive context-handling abilities. While versatile, GPT-4 needs enhancements for specialized tasks like TAA discovery. The Graph RAG approach [6] employs cutting-edge retrieval algorithms to gather relevant biological data, which GPT-4 then synthesizes. Leveraging advancements in NLP and data retrieval, the system generates detailed insights into TAA gene-cancer type connections, enhancing interpretative capabilities and information accuracy.

Our RAG framework is refined through continuous updates with new findings and iterative graph structure modifications, ensuring accurate TAA identification and characterization. This dynamic approach allows seamless integration of new data, keeping the model at the forefront of TAA research. A reinforcement component within the RAG framework implements a feedback loop, critically evaluating outputs to further refine the model. This continuous learning mechanism is crucial for maintaining performance and relevance in the evolving cancer research landscape.

We carefully integrate prompt templates, queries, and gene-aware contexts to construct comprehensive prompts. These prompts are submitted to GPT-4 with key parameters configured to facilitate the generation of nuanced completions. The Graph RAG-enhanced LLM output includes assessments of TAA gene-cancer type relationships and human-readable explanations, as well as safety assessment based on Omics data analysis, aiding scientific understanding and supporting therapeutic development with domain specific knowledge graph (e.g. **Fig. 1**). By refining and expanding the framework, we ensure the model remains a valuable tool for advancing cancer research and therapy innovation

**Figure 1:**
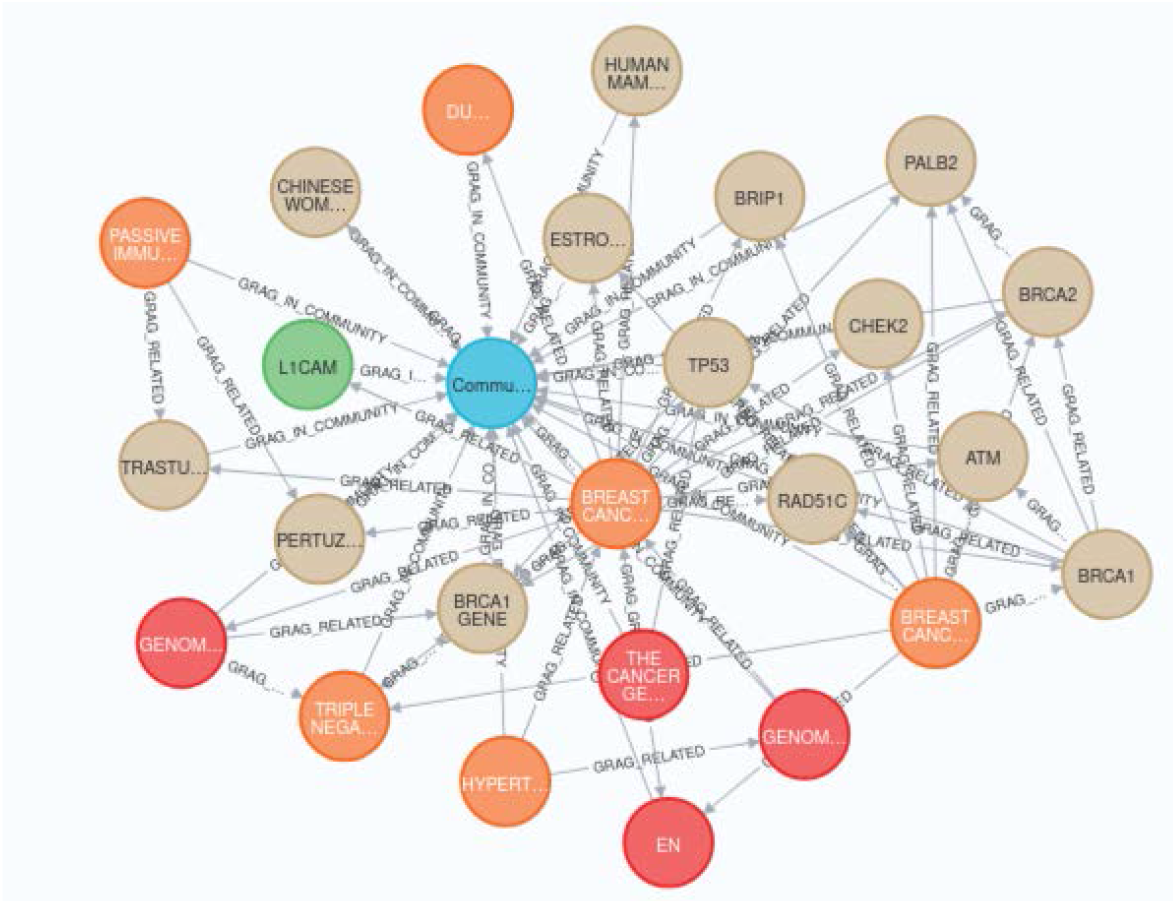
A subset of knowledge graph extracted from the given PubMed articles. Knowledge graph of entity nodes and relationship edges derived from the PubMed dataset (described in the txt), with different colors representing various communities.

## Results

An online service platform has been developed, allowing users to input prompts that include gene symbols and a cancer type to efficiently obtain automated results. These results indicate the significance of the gene(s) in the specified cancer type, accompanied by human-readable explanations and a summary of data-driven learning outcomes, integrated with omics data derived safety considerations. We assess the framework’s performance using known TAAs from clinical trials and compare the responses generated by the GraphRAG-enhanced LLM with the foundational model.

Here is an example comparing the responses to the query: “Evaluate TACSTD2 and MUC1 as Tumor-Associated Antigens in NSCLC; which one is a better ADC/TCE target?” between the outputs of the base model and the RAG-enhanced model:

**Table 1:**
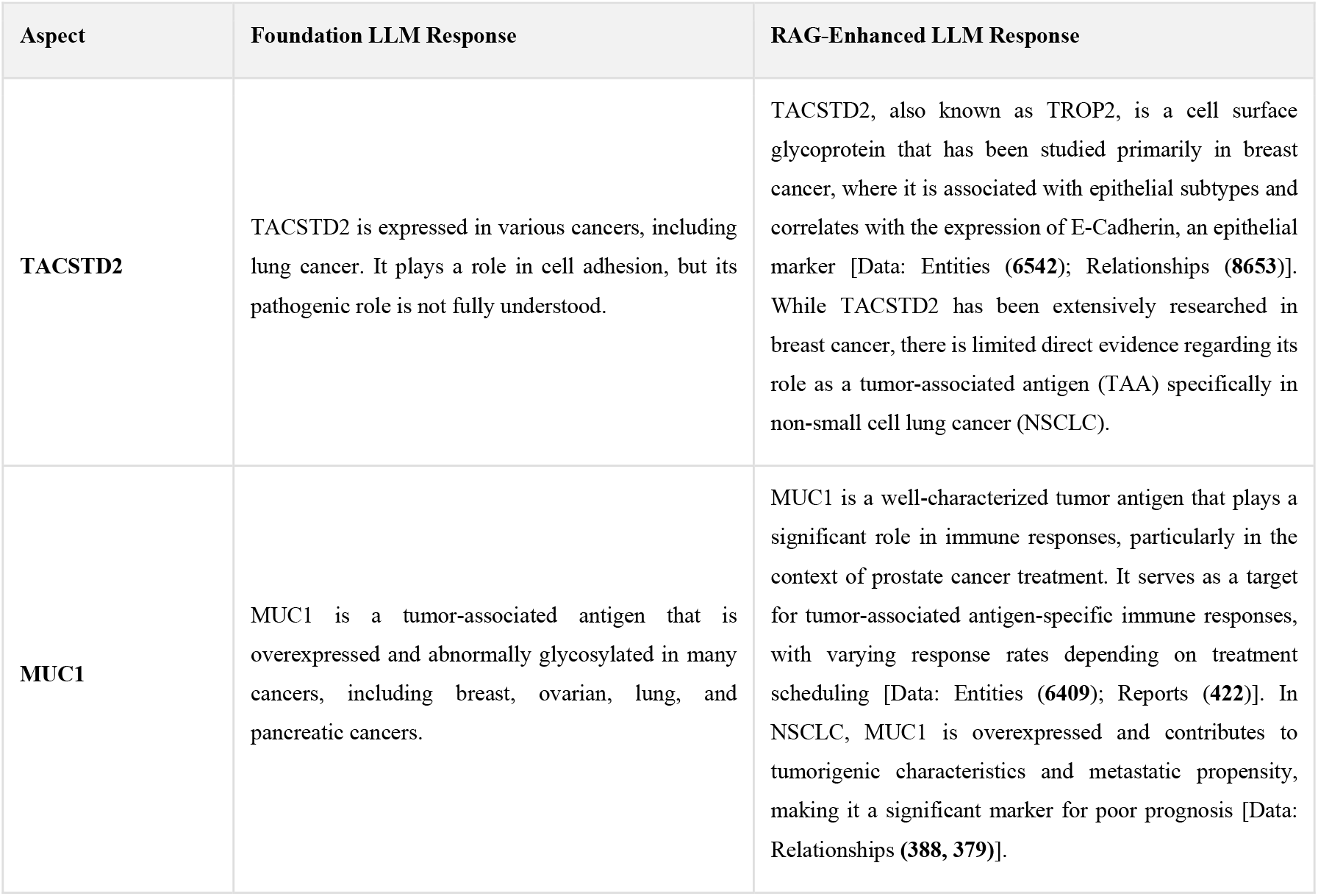
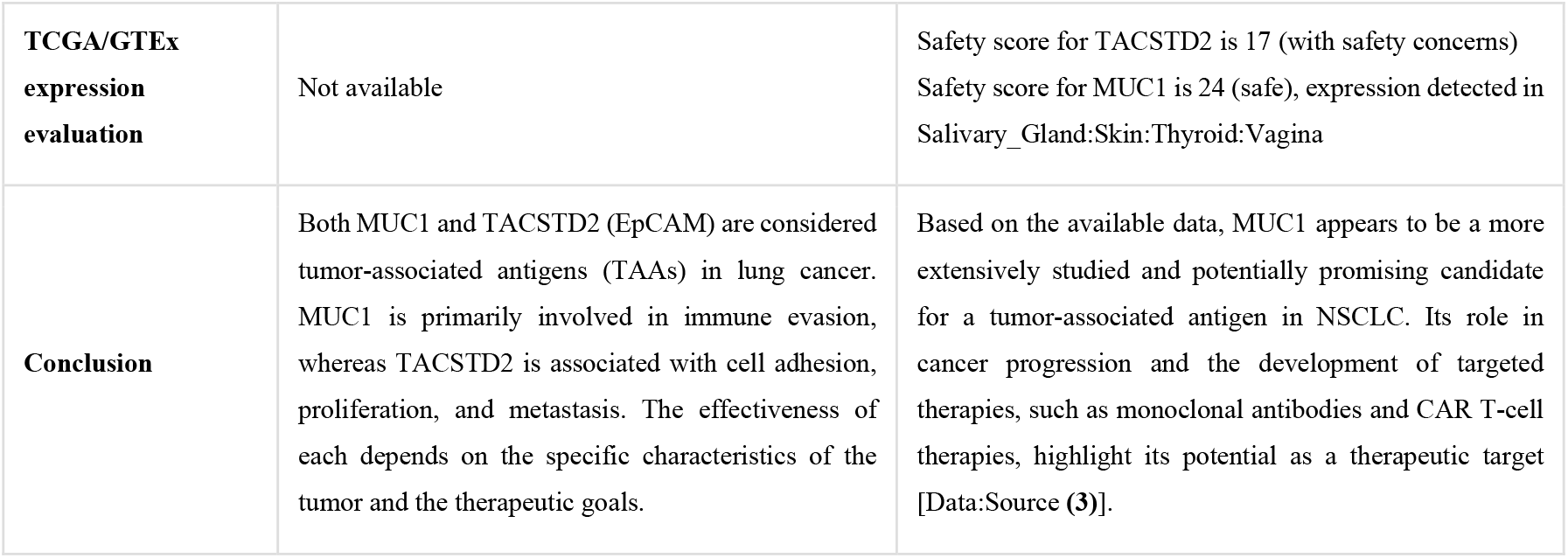
Model Response Comparison

| Aspect | Foundation LLM Response | RAG-Enhanced LLM Response |
| --- | --- | --- |
| TACSTD2 | TACSTD2 is expressed in various cancers, including lung cancer. It plays a role in cell adhesion, but its pathogenic role is not fully understood. | TACSTD2, also known as TROP2, is a cell surface glycoprotein that has been studied primarily in breast cancer, where it is associated with epithelial subtypes and correlates with the expression of E-Cadherin, an epithelial marker [Data: Entities (6542); Relationships (8653)]. While TACSTD2 has been extensively researched in breast cancer, there is limited direct evidence regarding its role as a tumor-associated antigen (TAA) specifically in non-small cell lung cancer (NSCLC). |
| MUC1 | MUC1 is a tumor-associated antigen that is overexpressed and abnormally glycosylated in many cancers, including breast, ovarian, lung, and pancreatic cancers. | MUC1 is a well-characterized tumor antigen that plays a significant role in immune responses, particularly in the context of prostate cancer treatment. It serves as a target for tumor-associated antigen-specific immune responses, with varying response rates depending on treatment scheduling [Data: Entities (6409); Reports (422)]. In NSCLC, MUC1 is overexpressed and contributes to tumorigenic characteristics and metastatic propensity, making it a significant marker for poor prognosis [Data: Relationships (388, 379)]. |
| **TCGA/GTEx expression evaluation** | Not available | Safety score for TACSTD2 is 17 (with safety concerns)<br>Safety score for MUC1 is 24 (safe), expression detected in Salivary_Gland:Skin:Thyroid:Vagina |
| **Conclusion** | Both MUC1 and TACSTD2 (EpCAM) are considered tumor-associated antigens (TAAs) in lung cancer. MUC1 is primarily involved in immune evasion, whereas TACSTD2 is associated with cell adhesion, proliferation, and metastasis. The effectiveness of each depends on the specific characteristics of the tumor and the therapeutic goals. | Based on the available data, MUC1 appears to be a more extensively studied and potentially promising candidate for a tumor-associated antigen in NSCLC. Its role in cancer progression and the development of targeted therapies, such as monoclonal antibodies and CAR T-cell therapies, highlight its potential as a therapeutic target [Data:Source (3)]. |

Based on responses from both models, we found that GraphRAG enhances LLM outputs with more relevant analysis for TAA candidates and uses engineered prompts to organize the output effectively. The conclusion that MUC1 is a better TAA for lung cancer aligns with current clinical observations, including its use in ADC and TCE therapies [7]. The foundation model is less specific and even wrongly connected TACSSTD2 (Trop2) to another common cancer marker EpCAM.

The enhanced LLM identified promising TAAs, particularly in response to ADC/TCE therapeutics that induce immune responses and alter cellular protein landscapes. Key candidates such as MUC1 demonstrated these antigens represent viable targets for ADC and TCE therapeutic development. Additionally, the integrated analysis has allowed detailed safety profiling of gene expression, providing deeper insights into contextual dynamics of antigen presentation in normal tissues. This enhances understanding beyond traditional methodologies, highlighting the potential for integrating LLMs with domain knowledge to guide targeted therapeutic approaches.

## Conclusion and Discussion

In cancer research, large language models are being used to analyze data and interact with patients, offering new perspectives on personalized treatment plans [4]. Our RAG-enhanced LLM offers a novel approach to TAA-based cancer therapy development, promising accelerated discovery processes and streamlined development pathways. Integration of LLMs with biological domain knowledge offers revolutionary potential in target discovery and prioritization, enhancing precision in cancer therapy development. Besides the TCGA/GTEx bulk RNAseq data for safety consideration, scRNAseq and proteomics data from cancer or normal samples can also be integrated. Our findings underline the importance of LLM’s potential and herald a new wave of effective, selective therapeutic strategies. However, despite the promising potential, it is crucial to undertake rigorous validation of the identified TAAs through in vivo studies. Additionally, exploring combinatorial drug approaches and continuously refining our LLM-enhanced platform could significantly boost predictive power and applicability across various cancer types, thereby guiding future research and applications. Enhanced collaboration among AI researchers and oncologists will be vital in harnessing the full potential of LLMs to transform cancer treatment landscapes. The source code for generating the TAA knowledge GraphRAG will be made available on GitHub.

In summary, this study presents a GraphRAG-enhanced model designed to identify and prioritize tumor-associated antigens (TAAs) for targeted cancer therapies. Using advanced NLP technologies, we demonstrated the potential of the enhanced LLM for TAA discovery and expedite therapeutic development for ADC and TCE therapies. Besides integrating omics features and rationales for creating detailed and interpretable analytical reports, it can be further extended to more biological datatypes such as image data from digital pathology [8]. The integration of the powerful LLM with the multimodal omics data-derived features presents a promising approach for improving the identification of cancer genes and elucidating their roles in cancer development and progression.

## Acknowledgments

We acknowledge the guidance of the development of the project by Jadwiga Bienkowska and s contributions of our collaborators and data providers.

